# Achilles tendon structure in distance runners does not change following a competitive season

**DOI:** 10.1101/324020

**Authors:** Todd J. Hullfish, Kenton L. Hagan, Ellen Casey, Josh R. Baxter

**Author notes:** Corresponding author: Josh R. Baxter, PhD. Mailing address: 3737 Market Street, Suite 1050, Philadelphia, PA, USA 19104. **Ethics Approval and Consent to Participate** this study was approved by the Institutional Review Board at the University of Pennsylvania (#827145). Subjects provided written-informed consent. **Consent to Publish** this submission does not contain any individual data. **Authors’ Contributions** All authors conceived the study; JB and EC designed the study, JB, TH, KH and EC acquired the data; JB, TH, and EC analyzed the data; JB and TH drafted the article; all authors revised the article critically for important intellectual content; all authors approved the submitted version.

## Abstract

Achilles tendon structure differs between trained distance runners and healthy controls, but the progression of tendon remodeling over the course of a competitive season is poorly understood. Therefore, the purpose of this study was to quantify Achilles tendon structure at the beginning and completion of a cross country season. We hypothesized that athletes who did not develop tendinopathy would not present with changes in tendon structure. Ultrasound assessments of the right Achilles tendon mid-substance were performed to quantify tendon organization, thickness, and echogenicity. Subjective structural measures and reported outcomes were also collected to determine if tendinopathy was present in any of the subjects. None of the subjects developed symptomatic tendinopathy over the course of the competitive season, but one runner did show signs of mild neovascularization. Tendon organization and echogenicity did not change over the course of the season. Tendon thickness increased by 7% (*P* < 0.001) but the effect size was small (d = 0.36). Runners who do not develop symptomatic tendinopathy have habituated tendon structure that may serve as a protective mechanism against the rigors of distance running. Monitoring tendon structure may serve as a means of detecting signs of structural indicators of tendinopathy prior to the presentation of symptoms.

## New & Noteworthy

Distance runners are at a 10-fold risk of developing Achilles tendinopathy, a painful degenerative condition. This work demonstrates that runners who do not develop symptoms over the course of a competitive-season have structurally stable tendons, suggesting a habituated tissue that can withstand the increased loads experienced during high-mileage running. Monitoring tendon structure in athletes may serve as a way to identify detrimental structural changes before symptoms become present.

## Introduction

Achilles tendinopathy is a painful degeneration of the tendon that is ten-times more common in running athletes compared to age-matched peers (14). During running, the Achilles tendon is cyclically loaded in excess of twelve body weights (13), which may be the driving factor in tendon remodeling and tendinopathy development in these athletes. However, the progression of load induced tendon remodeling in a population at increased risk of developing symptomatic tendinopathy is not well understood (23).

Structural changes associated with symptomatic tendinopathy such as decreased collagen alignment (12) – or ‘organization’ – and increased tendon thickness (19) have both been reported in asymptomatic athletic populations. Competitive collegiate distance runners have thicker and less organized tendons than their recreationally active peers (**Figure 1**), despite displaying no symptoms of tendinopathy (10). Similarly, hypertrophy of the Achilles tendon has been observed in both elite and recreational athletes (5, 20), suggesting structural adaptations in response to the cyclic loading experienced during running.

**Figure 1.**
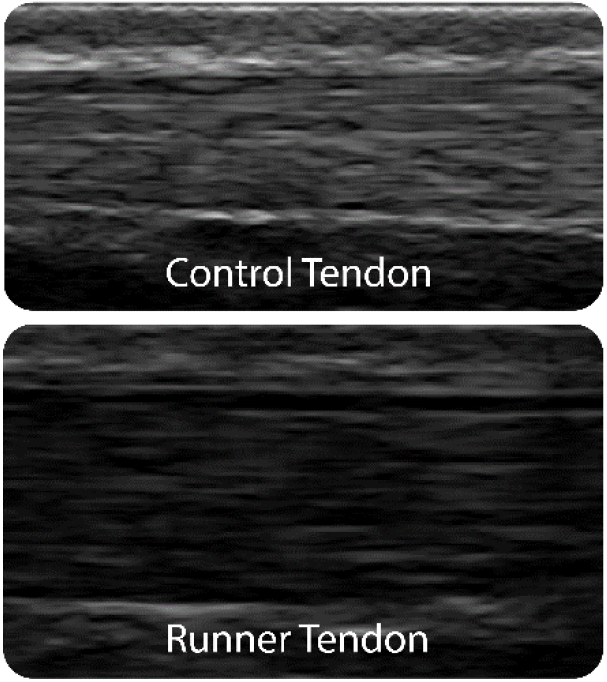
Major differences between the runner and control group can be seen when comparing these tendons under ultrasound. Runner tendon appears visibly thicker and less echogenic than control tendon. Reprinted with permission [pending] from *J Appl Physiol*.

Achilles tendon structure and properties change in response to training and previous exposure. Collagen synthesis in the peritendon increases following an intense bout of exercise (18) and prolonged periods of training in novice runners (17), which have not been associated with changes in tendon structure (8). Interestingly though, resistance training that involves high tendon strains stimulate stiffer tendons despite no increase in tendon cross-sectional area (3). Conversely, low strain magnitude training increases tendon cross-sectional area without increasing tendon stiffness (3). Trained runners have structurally different tendon compared to their recreationally active peers (10, 20, 26). However, it is unclear how tendon structure in highly-trained runners changes in response to prolonged bouts of training.

The purpose of this study was to prospectively quantify Achilles tendon structure in competitive distance runners at the beginning and completion of a cross country season. We hypothesized that, in the absence of injury, there would be no detectable differences in tendon thickness, organization, or echogenicity in the Achilles tendons of competitive distance runners. Should signs or symptoms of tendinopathy develop, we expected to detect structural changes associated with tendinopathy: increased thickness, decreased organization, and increased echogenicity (32). Understanding the structural remodeling response of tendon after exposure to prolonged bouts of cyclic loading is crucial to understanding how tendon disorders progress.

## Methods

### Study Design

Nineteen collegiate cross country runners (9 females; Age: 19 ± 1.5; Height: 172 ± 7 cm; Weight: 60.4 ± 8 kg; years running: 6.4 ± 2.5 yrs) provided written consent before participating in this IRB approved study. All participants had no symptoms of Achilles tendinopathy prior to participating in this study. Ultrasound images and patient reported outcomes were acquired one week prior to and one week following competing in a Division I NCAA Cross Country season. These visits took place on one of the team’s scheduled rest days and at least 12 hours after the last training session. Each study visit consisted of a self-reported assessment of tendon health and a quantitative ultrasound assessment. Participants completed a clinical-outcome questionnaire specific to Achilles tendon health (VISA-A)(11) in order to determine Achilles tendon healthy and function.

### Image Acquisition

Longitudinal B-mode ultrasound images of the mid-substance of the right Achilles tendon were acquired while subjects lay prone on a treatment table with ankles placed in the resting position off the end of the table. This position has been used previously to normalize the amount of tension in each subject’s tendon (30). Images were acquired with an 18 MHz transducer (L18-10L30H-4, SmartUs, TELEMED) with a 3 cm scanning width that was positioned approximately 4 cm proximal to the calcaneal insertion. Scanning parameters were held constant for all subjects over both scanning sessions (scan parameters: Dynamic Range: 72dB; frequency: 18 MHz; gain: 47 dB). Longitudinal Doppler images were acquired in order to quantify tendon neovascularization, which is a clinical indicator of tendon healing and pathology (32). Images were acquired by a single investigator as a continuous video and saved as video files.

### Image Analysis

Tendon structure was quantified using objective measures of organization and thickness. A single frame of each acquisition that clearly showed the mid-substance of the Achilles tendon (**Figure 2**) was selected by the same investigator for all subjects across both imaging sessions. In order to prevent any potential analyses bias, all images were selected and analyzed in a single session in which the trials were randomized and blinded to the investigator.

**Figure 2.**
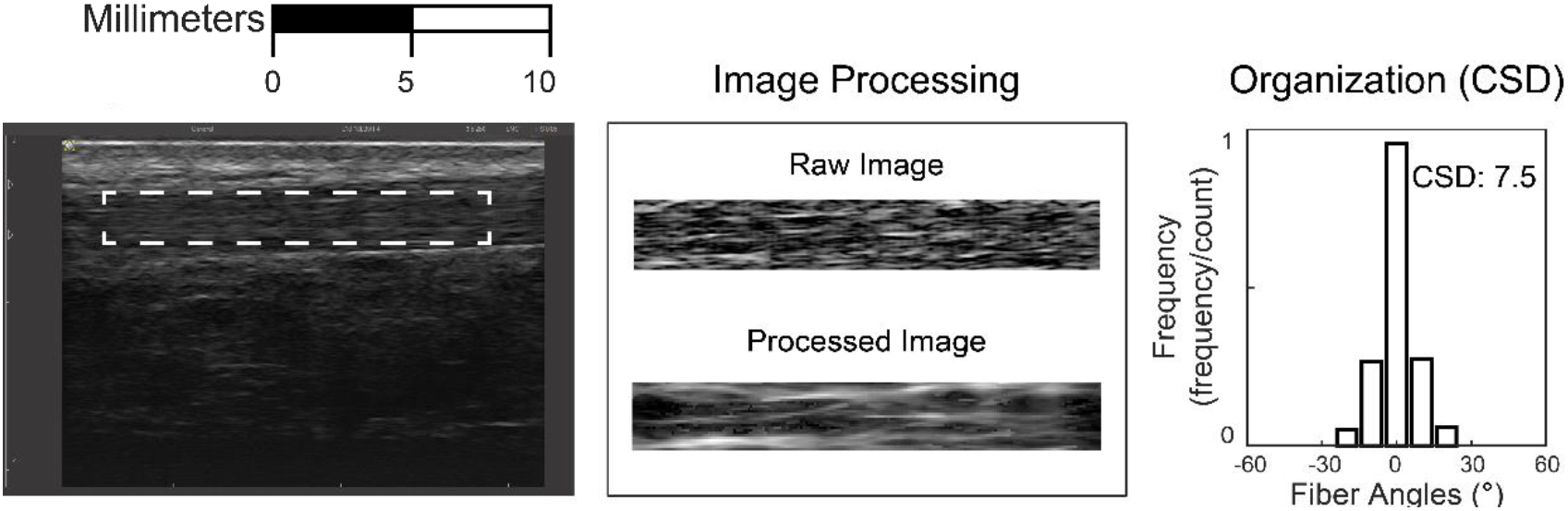
Rectangular regions of interest that encompassed as much of the free Achilles tendon as possible were independently selected by three investigators (*left*). Selections were processed computationally to identify the fascicles of the tendon (*middle*). Collagen fascicle alignment was calculated based on the deviation from the mean direction of all identified fascicles, termed the circular standard deviation (CSD). Reprinted with permission [pending] from *J Ultrasound Med*.

Collagen organization was quantified in the longitudinal B-mode ultrasound images using a computational analog for crossed polarizer light imaging (22). This analysis was implemented through a custom-written software that has been described in great depth elsewhere and has strong intrarater reliability (9). Briefly, the banded appearance of tendon results from collagen fascicles appearing hyperechoic and the non-collagenous matrix between the fascicles appearing hypoechoic under US (**Figure 2**, **middle**). To quantify the alignment of these bands, a rectangular region of interest of the visible length of the free Achilles tendon was selected (**Figure 2**, **left**), while edges of the tendon were avoided to prevent edge artifacts during analysis. A linear kernel was applied over groups of pixels in the image through a convolution matrix at varying angles between 0 and 180 degrees. The primary direction of each fascicle was calculated as the angle of maximum echo intensity using a power series function. The number of fascicles aligned at each angle was plotted in a histogram (**Figure 2**, **right**). The circular standard deviation (CSD) was calculated by determining the number of collagen fascicles that were not aligned in the primary fiber direction and the magnitude of the difference between their alignment and the mean angle. The CSD and collagen organization were then, by definition, inversely proportional: the higher the CSD, the less organized the structure of the tendon.

Tendon thickness and echogenicity were calculated using established-quantitative methods (6, 28, 32). Longitudinal thickness of the tendon was measured as the point to point distance between the deep and superficial edges of the tendon. Tendon thickness was calculated by the same blinded investigator using open-source image processing software (28). This measurement has been shown to be both repeatable and reliable in a prior report (32). Average echogenicity was calculated from the same regions of interest that were used to calculate collagen organization (6).

Clinical grading of tendon structure and neovascularization was performed by a fellowship-trained sports medicine physician who was blinded to all other subject data. Tendon structure and neovascularization were both graded (32) on a scale from 0 to 3: ranging from normal (0) to severe symptoms (3).

### Statistical Analysis

Tendon organization, thickness, and echogenicity as well as VISA-A scores were compared between the two study visits using two-way paired t-tests. Additionally, effect sizes were determined for any differences found to be statistically significant (P < 0.05). Effect sizes were reported using Cohen’s d, calculated as the mean difference divided by the pooled standard deviation. (27) Secondary regression analysis was performed to determine the relationship between organization and echogenicity.

## Results

Achilles tendon symptoms did not develop in any of the runners over the duration of the competitive running season, which was confirmed by clinical US exams and VISA-A scores at each imaging session (**Table 1**). During the first session, all runners were qualitatively determined to have ‘normal structure’. Prior to the competitive running season, one runner showed signs of ‘mild neovascularization’ (score of 1 out of 3) under Color Doppler, while all other runners had no signs of neovascularization. No changes were observed in clinical measures of tendon structure at the second session (**Table 1**). No detectable change in VISA-A scores between the pre- and post-season sessions (*P* > 0.1, Table 1).

**Table 1.** Subjective measurements of tendon status.

|  | Session 1 | Session 2 |
| --- | --- | --- |
| VISA-A (out of 100) | $93 \pm 8.1$ | $94 \pm 6.9$ |
| Structure (0 – 3) | $0 \pm 0$ | $0 \pm 0$ |
| Neovascularization (0 –3) | $0.05 \pm 0.21$ | $0 \pm 0$ |

Tendon organization and echogenicity did not change over the course of the competitive season (P > 0.05, **Figure 3**). Tendon thickness increased by 7% (P < 0.001, **Figure 3**), but the effect size of this change was small (d = 0.36). Mean echogenicity was negatively correlated with collagen alignment at both time points (Session 1: R^2^ = 0.24, *P* < 0.05; Session 2: R^2^ = 0.29, *P* < 0. 05, **Figure 4**).

**Figure 3.**
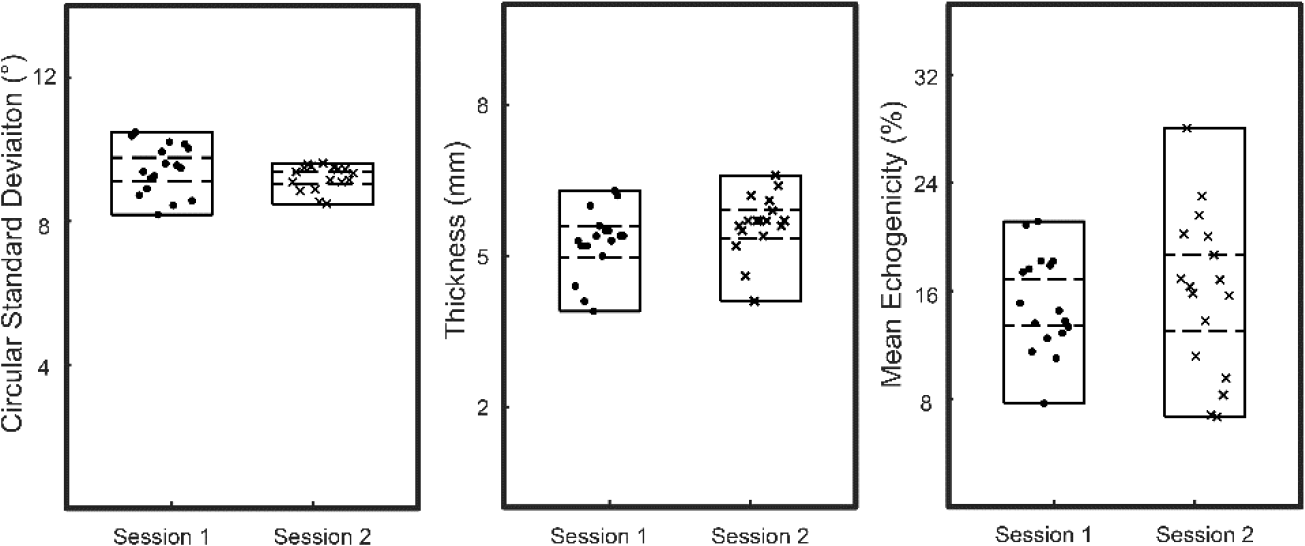
Measures of circular standard deviation (left), tendon thickness (center), and mean echogenicity (right) are plotted for preseason (session 1, N=19) and post-season (session 2, N=19). Measurement ranges (solid) and 95% confidence intervals (dashed) are plotted for each group. Circular standard deviation and mean echogenicity did not differ between sessions but tendon thickness showed a small increase following the training season (p < 0.001, effect size: 0.36).

**Figure 4.**
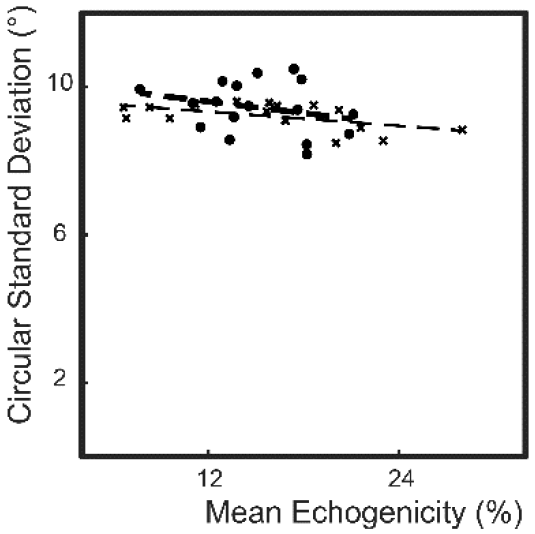
Mean echogenicity and circular standard deviation was negatively correlated before (session 1, R^2^ = 0.24, *P* < 0.05, dots) and following the competitive season (session 2, R^2^ = 0.29, *P* < 0.05, crosses).

## Discussion

Achilles tendon injuries are particularly common in running athletes (14), but the implications of a competitive training season on tendon structure is unclear. Therefore, the purpose of this study was to prospectively quantify Achilles tendon structure in competitive distance runners at the beginning and completion of a cross country season. We confirmed our hypothesis that competitive distance runners have Achilles tendon structure that is habituated to prolonged cyclic loading and does not change over a competitive season. While there was a statistically significant increase in tendon thickness (7%, P < .001) the effect size of this change was small (d = 0.36) and was not associated with any increases in clinical markers of tendinopathy (**Table 1**).

Our findings that tendon undergoes small amounts of hypertrophy throughout a competitive running season while maintaining a habituated structure is supported by previous literature. A previous ultrasound study on tendon thickness in distance runners also showed small amounts of tendon hypertrophy during a competitive season (31). Ultrasound analyses of shoulder tendons in elite swimmers shows darker and thicker tendons, suggestive of habituated tendon structure in response to long-term participation in training (25). Hypertrophy of the patellar and Achilles tendons have both been reported in recreational and elite athletes as a response to sport specific training (5, 8, 20).

Changes in tendon structure appear to be a protective adaptation (10, 20), mitigating injury risk in trained runners who cyclically load their tendons over extended periods of time. Mechanically, thicker tendon undergoes reduced strain when exposed to cyclic loading during running (26). In addition to thicker tendons, distance runners also have decreased collagen organization and echogenicity (10), which may further modulate the mechanical properties of the tendon (7). Tendon remodeling as a protective mechanism appears to be activity dependent, with sprinters having stiffer tendons than both distance runners and healthy controls (2). Despite similarities in tendon stiffness between distance runners and non-runners (2) the same study also demonstrated no difference in maximal torque generation potential between distance runners and non-active populations.

The intrinsic and extrinsic factors that drive tendon remodeling remain unclear but many mechanisms have been identified as potential drivers. Exercise has been shown to increase levels of collagen synthesis in humans (17, 18) but the effects of this increase have not been directly linked to tendon remodeling. These studies have been limited to short term changes in activity level, so it is unclear if increased collagen synthesis levels are maintained through long term training regimens or return to nominal levels following prolonged habitual training. The magnitude of tendon strain during training impacts tendon remodeling, both in terms of hypertrophy and increased stiffness (1). Additionally, lower frequencies of applied strain elicit larger changes of tendon mechanical properties (3). The varying mechanical adaptations elicited as a result of varying strain frequencies seem to agree with differences observed between long distance runners and sprinters (2). Distance runners are primarily interested in maintaining velocity and minimizing energy expenditure. In order to minimize energetics during running, trained-runners increase tendon loading in order to efficiently store and return elastic energy. Metabolic cost of distance running has been negatively correlated with Achilles tendon moment arm (21, 29), a mechanism that increases the amount of tendon tension during locomotion.

The processes by which tendon remodels from a naïve to a habituated – and from a healthy to a pathologic – state are still not well understood. Remodeling is not observed in novice runners over the course of 34 weeks (8); however, these runners only participated in low-intensity middle-distance running with the goal of reducing the likelihood of injury. Similarly, the present study demonstrates that the habituated tendons of experienced runners do not show signs of remodeling following prolonged training. While training results in increased metabolic activity and tendon remodeling in certain cases, there appears to be an intensity and frequency threshold that must be crossed in order to elicit the remodeling response. Rapid changes in activity level may increase the development of tendinopathy (4), suggesting that the loading thresholds for tendon remodeling and pathologic development depend on the history of training. This study cohort maintained a regimented training routine and did not develop tendinopathy despite being at a nearly ten-fold higher injury risk (14).

Several limitations should be considered when interpreting these results. Ultrasound measurements were made by a single investigator in a single data collection session but these measurements have been shown to be repeatable and reliable between and within investigators (9). All participants in this study were part of the same collegiate cross-country team and underwent similar training regimens, but each athlete’s weekly mileage varied and we did not control for factors such as diet, running mechanics, medication exposure, and genetics. Tendon mechanical properties were not measured in this study; however, prior reports have linked tendon mechanical properties with organization (15, 16) and echogenicity (6).

In conclusion, although the Achilles tendons in experienced distance runners are structurally different than controls – suggesting mechanical adaptation to running – they do not experience substantive changes in Achilles tendon structure following a competitive season in the absence of injury. These findings build confidence that some degree of remodeling may act as a protective adaptation. It remains unclear if changes to the structure of the tendon are predictive of the development of tendon dysfunction; however, intensive training for collegiate cross country competition does not appear to elicit detrimental changes in tendon structure in seasoned runners. Monitoring tendon structure in running athletes may identify subtle changes in tendon structure before symptoms present. Understanding the processes by which the tendon remodels in both healthy and pathologic ways is crucial to better treating and preventing tendon disorders.

## Acknowledgments

We would like to thank Annelise Slater and Neza Stefanic for assistance in data collection and Jamel Jones for assistance in subject recruitment.

## References

1 Arampatzis A, Karamanidis K, Albracht K. Adaptational responses of the human Achilles tendon by modulation of the applied cyclic strain magnitude. J Exp Biol 210: 2743–2753, 2007.

2 Arampatzis A, Karamanidis K, Morey-Klapsing G, De Monte G, Stafilidis S.Mechanical properties of the triceps surae tendon and aponeurosis in relation to intensity of sport activity. J Biomech 40: 1946–1952, 2007.

3 Arampatzis A, Peper A, Bierbaum S, Albracht K.Plasticity of human Achilles tendon mechanical and morphological properties in response to cyclic strain. J Biomech 43: 3073–3079, 2010.

4 Clement DB, Taunton JE, Smart GW. Achilles tendinitis and peritendinitis: Etiology and treatment. Am J Sports Med 12: 179–184, 1984.

5 Couppe C, Kongsgaard M, Aagaard P, Hansen P, Bojsen-Moller J, Kjaer M, Magnusson SP. Habitual loading results in tendon hypertrophy and increased stiffness of the human patellar tendon. J Appl Physiol 105: 805–810, 2008.

6 Crevier-Denoix N, Ruel Y, Dardillat C, Jerbi H, Sanaa M, Collobert-Laugier C, Ribot X, Denoix J-M, Pourcelot P. Correlations between mean echogenicity and material properties of normal and diseased equine superficial digital flexor tendons: an in vitro segmental approach. J Biomech 38: 2212–2220, 2005.

7 Gimbel JA, Van Kleunen JP, Mehta S, Perry SM, Williams GR, Soslowsky LJ. Supraspinatus tendon organizational and mechanical properties in a chronic rotator cuff tear animal model. J Biomech 37: 739–749, 2004.

8 Hansen P, Aagaard P, Kjaer M, Larsson B, Magnusson SP. Effect of habitual running on human Achilles tendon load-deformation properties and cross-sectional area. J Appl PhysiolBethesdaMd 1985 95: 2375–2380, 2003.

9 Hullfish TJ, Baxter JR. A reliable method to quantify tendon structure using B-mode ultrasonography. J. Ultrasound Med.

10 Hullfish TJ, Hagan KL, Casey E, Baxter JR. Achilles Tendon Structure Differs Between Runners And Non-Runners Despite No Clinical Signs Or Symptoms Of Mid-Substance Tendinopathy. J Appl Physiol.

11 Iversen JV, Bartels EM, Langberg H. the Victorian Institute of Sports Assessment - Achilles Questionnaire (Visa-a) – a Reliable Tool for Measuring Achilles Tendinopathy. Int J Sports Phys Ther 7: 76–84, 2012.

12 Khan KM, Cook JL, Bonar F, Harcourt P, Åstrom M. Histopathology of Common Tendinopathies. Sports Med 27: 393–408, 1999.

13 Komi PV. Relevance of in vivo force measurements to human biomechanics. J Biomech 23 Suppl 1: 23–34, 1990.

14 Kujala UM, Sarna S, Kaprio J. Cumulative Incidence of Achilles Tendon Rupture and Tendinopathy in Male Former Elite Athletes. Clin J Sport Med 15: 133, 2005.

15 Lake SP, Miller KS, Elliott DM, Soslowsky LJ. Effect of fiber distribution and realignment on the nonlinear and inhomogeneous mechanical properties of human supraspinatus tendon under longitudinal tensile loading. J Orthop Res 27: 1596–602, 2009.

16 Lake SP, Miller KS, Elliott DM, Soslowsky LJ. Tensile properties and fiber alignment of human supraspinatus tendon in the transverse direction demonstrate inhomogeneity, nonlinearity, and regional isotropy. J Biomech 43: 727–32, 2010.

17 Langberg H, Rosendal L, Kj\aer M. Training-induced changes in peritendinous type I collagen turnover determined by microdialysis in humans. J Physiol 534: 297–302, 2001.

18 Langberg H, Skovgaard D, Petersen LJ, Bulow J, Kj\a er M. Type I collagen synthesis and degradation in peritendinous tissue after exercise determined by microdialysis in humans. J Physiol 521: 299–306, 1999.

19 Maffulli N, Sharma P, Luscombe KL. Achilles tendinopathy: aetiology and management. JR SocMed 97: 472–476, 2004.

20 Magnusson SP, Kjaer M. Region-specific differences in Achilles tendon cross-sectional area in runners and non-runners. Eur J Appl Physiol 90: 549–553, 2003.

21 Raichlen D, Armstrong H, Lieberman D. Calcaneus length determines running economy: implications for endurance running performance in modern humans and Neanderthals. J. Hum. Evol. (2011). doi: 10.1016/jjhevol.2010.11.002.

22 Riggin CN, Sarver JJ, Freedman BR, Thomas SJ, Soslowsky LJ. Analysis of collagen organization in mouse achilles tendon using high-frequency ultrasound imaging. J Biomech Eng 136: 021–029, 2014.

23 Riley G. The pathogenesis of tendinopathy. A molecular perspective. Rheumatology 43: 131–142, 2004.

24 Robinson JM, Cook JL, Purdam C, Visentini PJ, Ross J, Maffulli N, Taunton JE, Khan KM. The VISA-A questionnaire: a valid and reliable index of the clinical severity of Achilles tendinopathy. Br J Sports Med 35: 335–41, 2001.

25 Rodeo SA, Nguyen JT, Cavanaugh JT, Patel Y, Adler RS. Clinical and Ultrasonographic Evaluations of the Shoulders of Elite Swimmers. Am J Sports Med 44: 3214–3221, 2016.

26 Rosager S, Aagaard P, Dyhre-Poulsen P, Neergaard K, Kjaer M, Magnusson SP. Load-displacement properties of the human triceps surae aponeurosis and tendon in runners and non-runners. Scand J Med Sci Sports 12: 90–98, 2002.

27 Sawilowsky SS. New Effect Size Rules of Thumb. J Mod Appl Stat Methods 8: 597–599, 2009.

28 Schneider CA, Rasband WS, Eliceiri KW. NIH Image to ImageJ: 25 years of Image Analysis. Nat Methods 9: 671–675, 2012.

29 Scholz MN, Bobbert MF, van Soest AJ, Clark JR, van Heerden J. Running biomechanics: shorter heels, better economy. J Exp Biol 211: 3266–3271, 2008.

30 Silbernagel KG, Steele R, Manal K. Deficits in Heel-Rise Height and Achilles Tendon Elongation Occur in Patients Recovering From an Achilles Tendon Rupture. Am J Sports Med 40: 1564–1571, 2012.

31 Sponbeck J, Perkins C, Berg M, Rigby J. Achilles Tendon Cross Sectional Area Changes Over a Division 1 NCAA Cross Country Season. Int JExerc Sci 10: 1226–1234, 2017.

32 Sunding K, Fahlström M, Werner S, Forssblad M, Willberg L. Evaluation of Achilles and patellar tendinopathy with greyscale ultrasound and colour Doppler: using a four-grade scale. Knee Surg. Sports Traumatol. Arthrosc. (2014). doi: 10.1007/s00167-014-3270-4.

